# Rapid isolation of functionally intact nuclei from the yeast *Saccharomyces*

**DOI:** 10.1101/162388

**Authors:** Mario Niepel, Julia C. Farr, Michael P. Rout, Caterina Strambio-De-Castillia

**Author notes:** These authors contributed equally (Mario Niepel, Julia C. Farr). Corresponding authors: C.S.-D.-C. and M.P.R.

## Abstract

Most available methods for nuclear isolation entail lengthy procedures that are difficult to master and generally emphasize yield and enrichment over nuclear preservation, thus limiting their utility for further studies. Here we demonstrate a novel and robust method to rapidly isolate well-preserved yeast nuclei. The method can be easily adapted to multiple preparation scales depending on experimental need and it can readily be performed on multiple samples by a single researcher in one day. We show that the nuclei fraction is strongly enriched and that the resulting nuclei are free from contaminating endoplasmatic reticulum and other cell debris. EM studies show that preservation of nuclear morphology is exquisite, making it possible to study peripheral nuclear pore components such as the cytoplasmic filaments and the basket, whose structure is generally difficult to maintain ex vivo. In addition, incubation of isolated nuclei with bulk transport substrates of different sizes and with import cargo indicates that the nuclear envelope is intact and nuclear pores retain their capacity to bind transport substrates. Our results suggest that this preparation procedure will greatly facilitate studies of the yeast nucleus which have been difficult to establish and to multiplex to date.

## Introduction

For decades the isolation of enriched nuclear fractions and their use for subsequent experimental analysis has been invaluable for understanding the composition, structure, and function of nuclei and of their content. Pioneering work has focused on the isolation of nuclei from vertebrates (Dounce and Lan, 1943; Dounce, 1944; Blobel and Potter, 1966). The first methods to isolate nuclei from yeast, a key unicellular eukaryotic model organism, were reported for *Saccharomyces uvarum* (Rozijn and Tonino, 1964) and *Schizosaccharomyces pombe* (Duffus, 1969). Throughout the years, these methods have been adapted to work with the more commonly used *S. cerevisiae* strains (Bhargava and Halvorson, 1971; Wintersberger *et al.*, 1973; Hurt *et al.*, 1988; Aris and Blobel, 1991) and have provided the basis for the production of highly-enriched spindle pole bodies (Rout and Kilmartin, 1990), enriched nucleoli(Aris and Blobel, 1991), highly enriched nuclear pore complexes (NPCs; (Rout and Blobel, 1993)), and enriched nuclear envelopes (NEs; (Strambio-de-Castillia *et al.*, 1995)). Yet, except for notable exceptions (Strambio-de-Castillia *et al.*, 1995), these methods generally do not yield nuclei that are enzymatically active and competent for nuclear transport. This could be due to protein loss during isolation (Paine *et al.*, 1983) leading to damaged NPCs or to damage in the structural integrity of the NE during preparation (Blobel and Potter, 1966).

More recently, Beck and coworkers isolated *Dictyostelium discoideum* nuclei and demonstrated their import competence by providing an energy regenerating system and monitoring the import of fluorescently labeled BSA fused to a canonical nuclear localization signal (Beck *et al.*, 2004). Transport competent nuclei can also be obtained from *Xenopus laevis* oocytes by manual dissection (Evans and Kay, 1991) and such preparations have been used for studies of nucleocytoplasmic transport mechanism in mammalian cells (Keminer and Peters, 1999). In addition, in mammalian cells and *Xenopus laevis* oocytes, nuclear transport can also be explored by less invasive methods such as the microinjection of transport substrates directly into the cytoplasm to study nuclear import (Feldherr and Feldherr, 1960) or into the nuclei to study nuclear export (Gurdon *et al.*, 1976). Finally, the use of digitonin to permeabilize the plasma membrane of the target cells while leaving the NE and NPC intact, has provided the opportunity to monitor the process of nuclear import ex vivo (Adam *et al.*, 1992) leading in turn to the characterization of many essential soluble factors involved in the nuclear transport pathway (Sweitzer and Hanover; Takizawa *et al.*; Plafker and Macara). Unfortunately, such techniques are not available in yeast due to the presence of the cell wall, which limits access to the cellular interior. Therefore, in yeast the isolation of functionally and structurally intact nuclei is a necessary first step for the development of reconstituted nuclear transport systems.

Here we describe a new method to rapidly isolate functional yeast nuclei, which is a modification of previously described methods used in our lab (Rozijn and Tonino, 1964; Rout and Kilmartin, 1990; Rout and Blobel, 1993; Strambio-de-Castillia *et al.*, 1995; Kipper *et al.*, 2002) with the following advantages: 1) The method described here is much faster than previous procedures, allowing the production of enriched nuclei in parallel from multiple samples in approximately half a workday. 2) It yields structurally and functionally intact nuclei. 3) The method is easily scalable, allowing for the production of both small- and large-scale preparation, depending on experimental needs. 4) Nuclear fractions can be easily stored in small aliquots at -80C without apparent loss in transport efficiency or structural integrity of both NE and NPCs, further increasing multiplexing opportunities. By using a strain expressing a fluorescently tagged variant of Nup49p as the source of enriched nuclei, we were able to quickly monitor the procedure during isolation, which enabled us to optimize the process. Our method to quickly isolate nuclei from both wild type and mutant strains can thus form the basis of both structural and functional investigations of yeast nuclei.

## Results and Discussion

### Quick and easy isolation of yeast nuclei

Our new method to rapidly isolate highly enriched fractions of functionally intact nuclei in parallel from multiple sample types is outlined in Fig. 1. Yeast cells of the desired strains are grown to mid log-phase (1 × 10^7^ cells/ml for diploid and 2 × 10^7^ cells/ml for haploid strains) to prevent thickening of the cell wall, which can occur in late log- or stationary-phase. Cells are harvested by centrifugation, washed once in water, and the resulting pellet is disaggregated in 100 mM Tris pH 9.4, 10 mM DTT. The cell wall is removed by enzymatic digestion in the presence of 1.1 M sorbitol to avoid premature lysis of the resulting spheroplasts (Rout and Blobel, 1993; Strambio-de-Castillia *et al.*, 1995; Kipper *et al.*, 2002). Crucially, during digestion the preparation is subjected to periodic shaking, which is vigorous enough to keep yeast cells from clumping and sinking to the bottom, yet gentle enough to prevent damaging the resulting spheroplasts. The progress of the digestion can be monitored by comparing the appearance of the cells after dilution 1:20 in 1.1 M sorbitol versus water by phase contrast microscopy. In sorbitol, the spheroplasts should appear phase bright and spherical with no buds attached, while in water they should completely lyse with only fragments of the membranes visible. When digestion is complete, the spheroplasts are washed once and then pelleted through a cushion of 7.5 % Ficoll / 1.1 M sorbitol. Cells are lysed and intact nuclei are released from cellular debris with a polytron homogenizer in lysis buffer containing a small amount of Triton-X100, which facilitates membrane breakdown, and 8% polyvinylpyrrolidone (PVP), which stabilizes the nuclei. The progress of the lysis can be checked in between polytroning steps by phase contrast microscopy. Free nuclei appear phase dark and, at high magnification, nucleoli can be seen as darker areas. Vacuoles appear phase bright and should disappear completely. Lysis is judged completed when less than 2% of unbroken cells can be seen. The lysate is diluted in 8% PVP solution and the suspension is homogenized by an additional polytroning step, which reduces clumping of nuclei and helps to remove endoplasmic reticulum from the nuclear membrane. After homogenization the lysate is loaded on a two-step sucrose gradient (1.875 M and 2.5 M in BT Mg buffer) after gently softening the interface with a glass rod. Unlike previously described procedures, our method requires only a one-step gradient that additionally has a large concentration differential. This makes the gradient extremely easy to prepare and is robust enough so that concentration adjustment by refractive index are not needed. Nuclei are unable to migrate into the 2.5M sucrose layer, while the loading solution and contaminating soluble and membranous material get retained by the 1.875 M sucrose layer. This gradient composition produced high yields, while at the same emphasizing the degree of enrichment. Optionally, at this stage the isolated nuclear fraction can be subjected to further treatment with DNase I. This removes the bulk of the DNA and gives rise to DNA-extracted nuclear shells [Fig. S1]. Both highly enriched isolated nuclei and nuclear shells can be quickly frozen by dripping directly into liquid nitrogen where they form beads of 20-30 μl each. The beads can be stored at -80°C and individual beads can be retrieved and thawed for future experiments.

**Figure 1:**
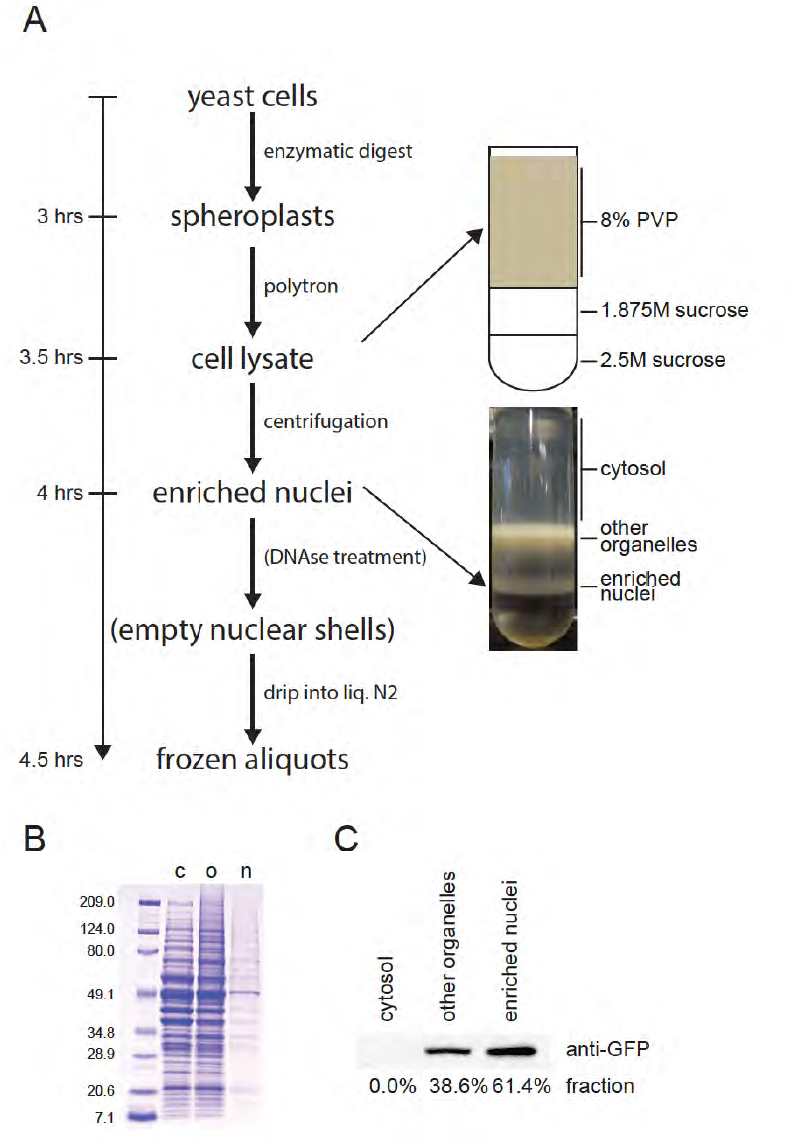
High-yield, rapid isolation of nuclei from the budding yeast. **(A)** Schematic depiction of the nuclei preparation procedure including images of the sucrose gradient loaded with lysate obtained from a total of 5×10^9^ spheroplasts (i.e. 0.5 l of media containing 1×10^7^ cells/ml), before and after centrifugation. Labels on the right show the composition of the different gradient steps (top) and the predominant content of the interfaces between layers (bottom). **(B)** Coomassie-stained SDS-PAGE profile of proteins found in fractions corresponding to the boundary regions identified in A. **(C)** Immunoblot of the proteins from the three boundary regions identified in (A) as revealed by staining with anti-GFP antibody.

### Nuclei are highly enriched and produced at high yield

To effectively monitor the purification process, we prepared nuclei from a *S. cerevisiae* strain expressing a Nup49p-GFP fusion protein (Ghaemmaghami *et al.*; Howson *et al.*). Nup49p is a core component of the NPC and it localizes almost exclusively to the NPC (Bucci and Wente). This makes the Nup49p-GFP strain ideal to follow the nuclear purification visually by fluorescence microscopy and biochemically by immunoblotting.

We prepared crude nuclei from the Nup49p-GFP strain and subjected them to gradient centrifugation as described above. After centrifugation we were able to distinguish three distinct layers by both macroscopic and microscopic visual inspection (photo in Fig. 1A). The first layer was clear and contained soluble cytoplasmic proteins (cytosol/c). The next layer was yellowish and harbored predominantly membranes bound organelles, plasma membrane remnants and other cellular debris (other organelles/o). The third, whitish layer, which was found at the 1.875 M – 2.5 M sucrose interface, was highly enriched in nuclei (nuclei/n). Microscopic observation also revealed that a yellowish color generally indicates contamination with dead cells, while the presence of a white smear between the nuclei and supernatant layer is due to broken nuclear material. We analyzed the protein content of each fraction by loading proportional volumes on SDS-PAGE. Protein staining by Coomassie showed a striking disparity in the protein concentration of the different fractions (Fig. 1B). While the cytosol and the other organelle fractions where rich in protein and contained similar major protein bands, the nuclei fraction contained relatively few proteins and showed a significantly different band pattern. Quantitative immunoblotting for Nup49p-GFP confirmed that even though the protein concentration in the nuclei fraction is low, it contains the majority of NE associated proteins (Fig. 1C). This indicates that as expected the nuclei fraction is highly enriched for components of the nucleus, such as the NPC.

Inspection by DIC microscopy (Fig. 2A) and fluorescence microscopy (Fig. 2B) confirmed that the nuclei fraction was highly enriched in spherical nuclei and that it was largely free of other cellular components or debris. Higher magnification showed that Nup49p-GFP maintained the distinct punctate rim-staining pattern characteristic of intact NPCs, suggesting that the gross structure of the NE was not disrupted (Fig. 2C). Further, to assess the presence of membranous contaminants in the nuclei fraction, we co-stained Nup49-GFP containing nuclei (Fig. 2D) with a lipid dye. The merged image shows that lipids appeared to localize exclusively to the nuclear envelope, as indicated by the presence of Nup49p-GFP. This is a good indicator that our procedure to release nuclei from spheroplasts removed all detectable contaminating ER remnants, which otherwise would radiate out from the nucleus leading to lipid stain independent of the GFP signal. In addition, the continuous circular pattern we observed with the lipid dye further underscored the degree of preservation of the nuclear envelope, and indicated that isolation procedure is rapid and gentle enough to preserve the gross structure of the nucleus.

**Figure 2:**
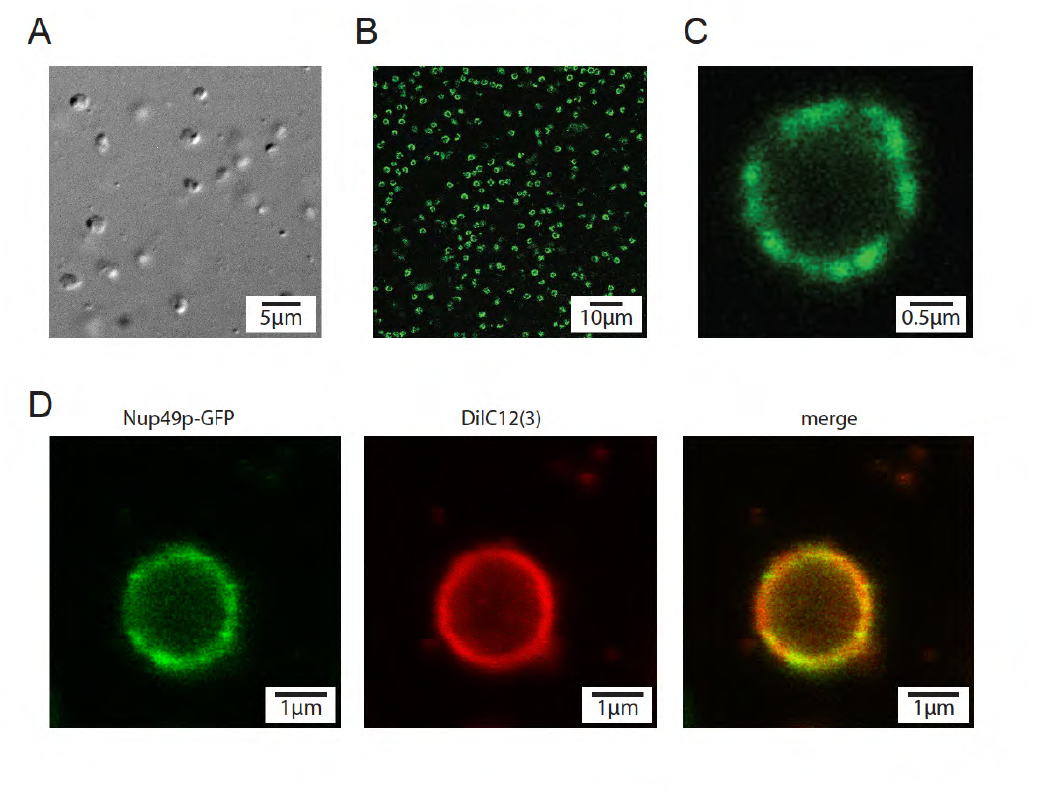
Nuclear fractions are highly enriched and show undetectable levels of membranous contaminants. **(A)** Bright-field microscopic image of the enriched nuclear fraction obtained from yeast cells expressing Nup49-GFP. **(B, C)** Fluorescence microscopy images of enriched nuclei obtained from cells expressing Nup49-GFP shown at **(B)** low and **(C)** high magnification **(D)** Florescence microscopy images of an individual representative nucleus showing signal associated with Nup49p-GFP, the membrane-dye DiIC12(3), and overlay of the two (merge), as indicated.

### Nuclei maintain structural integrity

To gain a more detailed view of the preservation condition of isolated nuclei, we observed them by electron microscopy (EM). In intact cells it is challenging to find fixation and staining methods that deliver contrast of both the nucleus and the cell, since yeast have dense and morphologically relatively undifferentiated protoplasm. However, freeing the nuclei from the surrounding cellular material results in superior morphological resolution, as shown previously (Rout and Kilmartin, 1990; Strambio-de-Castillia *et al.*, 1995)). Both low- and high- resolution images of enriched nuclei preparations reveal remarkable levels of structural and morphological preservation (Fig. 3A and B), with only a minority of the nuclei in any given field appearing damaged. At low magnification, nuclei appear round and their contours well preserved. At higher magnification, it appeared that the level of preservation of individual nuclear structures surpassed previous observations (Fig. 3B). Of particular interest was that when NPCs were sectioned through their mid-planes, we were able to clearly identify nuclear baskets (Fig. 3B, inset; (Niepel *et al.*)), which has been rarely observed in previously described nuclear preparations(Rout and Kilmartin, 1990; Fahrenkrog *et al.*, 1998; Kiseleva *et al.*, 2004) and virtually impossible to see in thin-sectioned whole cells. For a more precise investigation of the NE, scanning electron microscopy (SEM) was performed (Fig. 3C), which is a much gentler method to investigate nuclei by electron microscopy. Virtually all nuclei recognizable as such appeared as round spheres, free of ruptures in the NE. The exceptional preservation of sub-nuclear structures and the excellent condition of the NE seen by scanning EM further indicates that our method is very gentle and compatible with functional studies on isolated nuclei.

**Figure 3:**
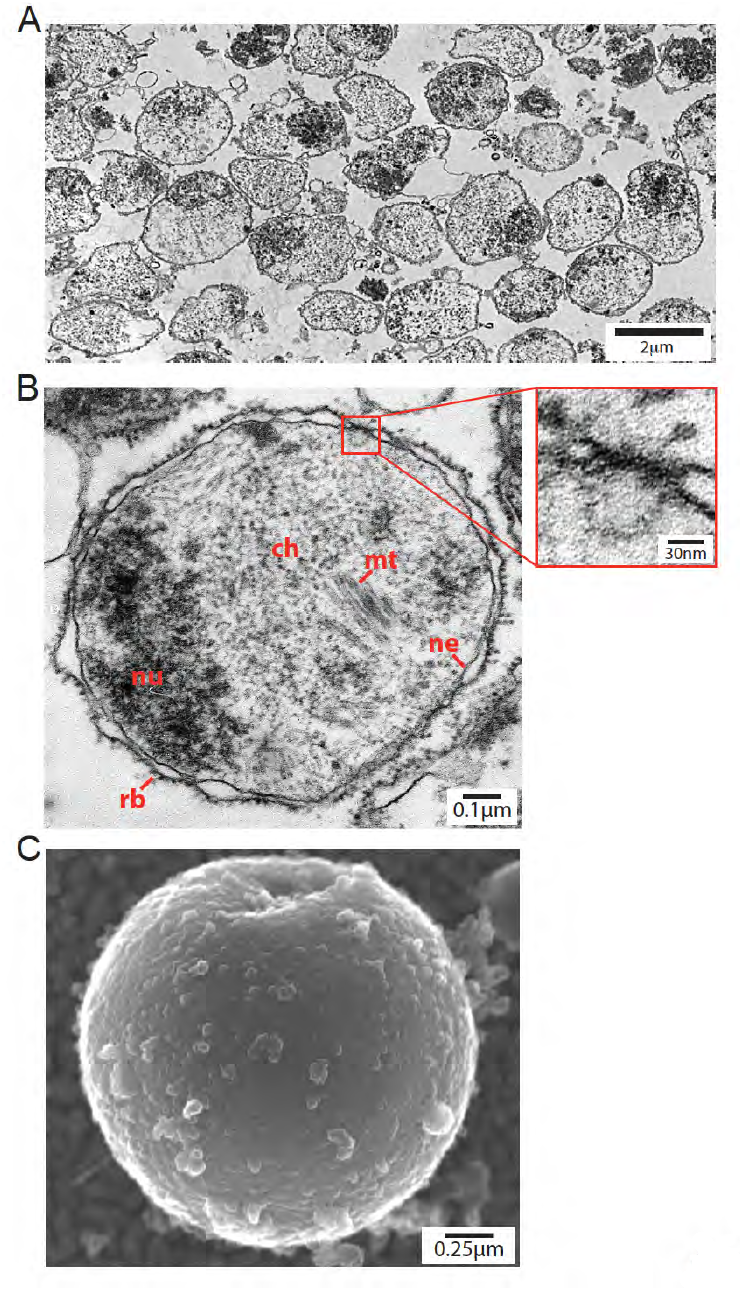
Electron microscopy images of isolated yeast nuclei show remarkable levels of morphological preservation. **(A)** Low magnification images showing prevalently well preserved nuclei with occasional clustered or broken nuclei. **(B)** High magnification image of a representative nucleus from the same section revealing the following structural details, the electron-dense structure on the left of the field constitutes the nucleolar compartment (nu), fuzzy structures found in the nucleoplasm constitute chromatin (ch), tubular structures represent transversal sections of microtubules (mt), and small dots decorating the surface of the outer nuclear membrane represent ribosomes (rb). Finally, the double membrane of the NE (ne) is visible around the whole nucleus. Inset: An enlarged section of the same nucleus is shown displaying a NPC, the nuclear basket reaching into the nucleoplasm, and the NE double membrane found on both sides of the NPC **(C)** As revealed by SEM, the observed nucleus is largely round and reveals an intact NE.

### Nuclear membrane is impermeable to large fluorescent cargo

We sought to further assess the structural and functional integrity of NEs and of NPCs after isolation. It has been shown that inert cargo larger than approximately 40 kDa or with a radius larger than 5 nm is not able to penetrate the NE or translocate through intact NPCs (Paine, 1975; Paine *et al.*, 1975; Peters, 1984). Consistently, both GFP (27 kDa) and BSA (67 kDa) have been shown to display very low transport rates of 3 molecules NPC-1 s-1 μm-1 (Ribbeck and Görlich, 2001) and 0.1 molecules NPC-1 s-1 μm-1 (Siebrasse and Peters, 2002) respectively. We found that if we add large proteins such as BSA labeled with Alexa-488 (67 kDa; Fig. 4A) or the naturally fluorescent phycoerythrin protein (240 kDa; Fig. 4B) to our fraction of purified nuclei, these proteins remain excluded from the nuclei (merge). While the area around nuclei showed bright fluorescence associated with the protein reporters, the nuclear interior, counter-stained for DNA, remained dark. However, nuclei did allow for slow entry of smaller proteins like Ovalbumin (46 kDa; Fig. 4C), which likely leaked into the nuclei by slow diffusion through the NPC. Remarkably, Ovalbumin was found to be mostly excluded from a crescent shaped area inside the nucleus, which appears to correspond to the nucleolus, as indicated by the lack of DRAQ5 counterstaining. In addition, the concentration inside the nucleus was significantly lower than outside indicating that the NPC barrier was still effective in preventing free diffusion in an out of the nucleus. In contrast, if the nuclear membrane were compromised it should have also allowed larger substrates to enter the nucleus and there should have been a rapid equilibration of the protein concentration inside and outside of the nucleus. We therefore can conclude that the nuclei are largely intact, and contain intact NPCs that effectively block diffusion of large molecules.

**Figure 4:**
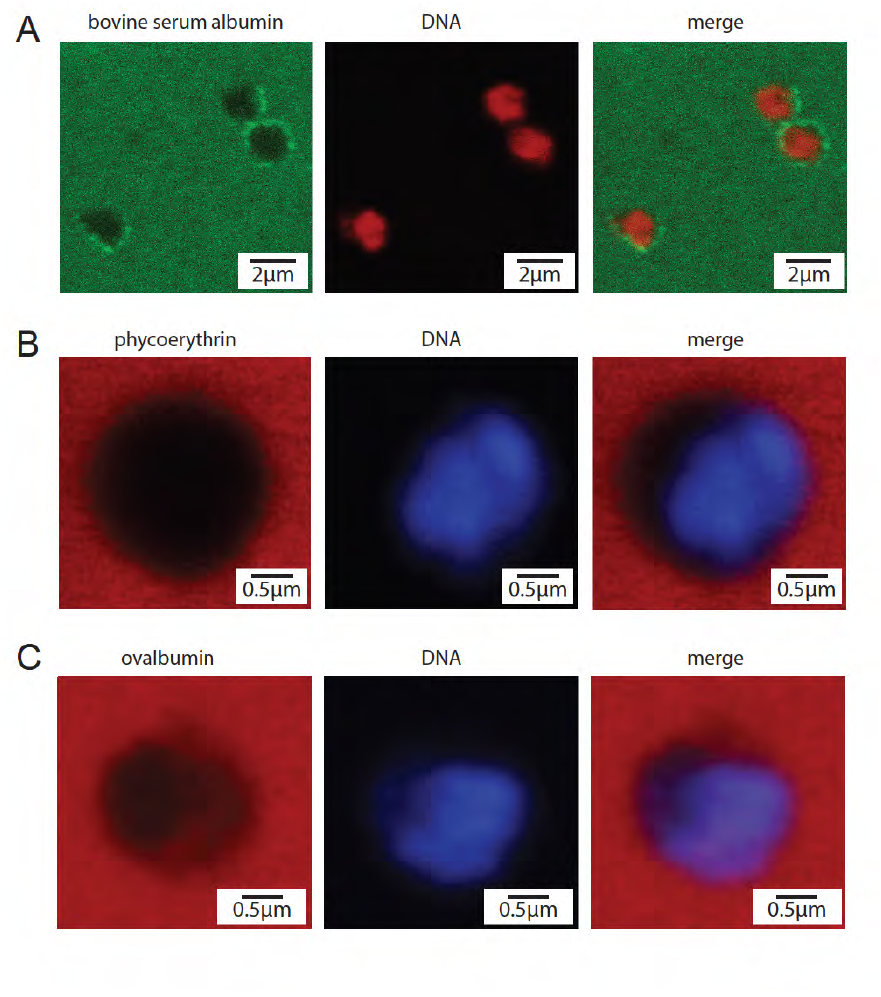
NE is impermeable to large fluorescent dyes. Isolated, Nup49-GFP tagged nuclei were incubated with 1 μM BSA-Alexa 488 (A), 250 nM R-phycoerythrin (B) or 1 μM ovalbumin-AlexaFluor-568 (C), and counterstained with 5 μM of DRAQ5 to reveal the position of chromosomal DNA before microscopic observation. **(A)** Green background fluorescence surrounding the nuclei is due to excess BSA-AlexaFluor-488, while the punctuate rim staining represents the localization of Nup49-GFP to the NE of isolated nuclei. No green signal is found inside the nuclei indicating that the NE barrier is preserved during nuclei preparation. **(B)** Red background fluorescence surrounding the nuclei is due to excess R-phycoerythrin (R-PE). No R-PE signal is found within the nuclei supporting the notion that the NE barrier remains intact after preparation. **(C)** Red background fluorescence surrounding the nuclei represents excess Ovalbumin, which is excluded from the nuclear interior. It should be noticed that while some Ovalbumin is able to diffuse to the nucleoplasm the majority is excluded. In addition, the nucleolus appears to remain mostly devoid of both DRAQ5 and of Ovalbumin-associated fluorescent signal.

### NPCs remain capable of binding transport substrates

To determine whether the isolated yeast nuclei retain transport competence and possess functional NPCs, we added Kap95p, the Importin β homolog in yeast, labeled with the fluorescent dye Cy5 (Seedorf and Silver, 1997) to the nuclear fraction labeled with Nup49p-GFP (Fig. 5). As seen earlier, Nup49p-GFP showed the characteristic punctate rim staining of NPCs. Localization of Kap95p-Cy5 was nearly indistinguishable from that of the NPCs; it showed a very similar punctate staining, which overlapped well in many of the larger NPC clusters (Fig. 5, merge). This indicated that Kap95p, in contrast to the transport-inert proteins observed in Fig. 4, bound specifically to NPCs. These findings strongly suggest that nuclei isolated by the method described here are competent for nuclear transport and contain intact NPCs.

**Figure 5:**
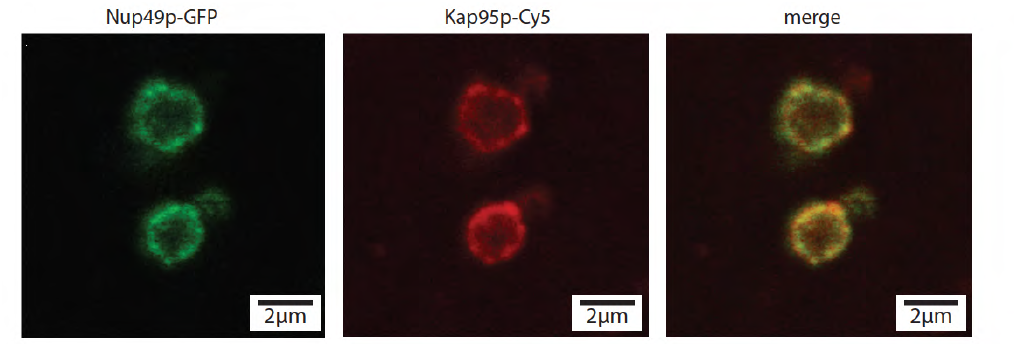
NPCs are competent for karyopherin binding. Fluorescence microscopy images of Nup49p-GFP nuclei treated with Kap95p-Cy5. Fluorescence of Nup49p-GFP (left panel), Kap59p-Cy5 (middle panel), and a merged image (right panel). All images have been smoothed using Gaussian blur (radius of 1 pixel).

## Conclusions

The rapid method for isolating yeast nuclei from both haploid and diploid *Saccharomyces* strains described here has key advantages over those available in the literature. High quality nuclei isolation procedures described to date, including those by our laboratory, normally require lengthy preparation and execution times, including multiple differential centrifugation steps and extensive sample manipulation (Bhargava and Halvorson, 1971; Aris and Blobel, 1991; Kipper *et al.*, 2002). In contrast, the method described here yields intact and highly enriched nuclei directly from whole cell lysates by performing a single ultracentrifugation run of less than ½ hour. This breakthrough dramatically cuts the time required to carry out the procedure and sample manipulation reducing total duration to half a day of work. At the same the procedure is inherently robust because the simplicity of the two-step sucrose gradient, characterized by a high sucrose concentration differential, eliminates the requirement for careful refractive index adjustments. Reduced sample handling and the robust gradient also significantly decrease the learning curve of the procedure and the possibility for variance in the quality of the nuclei from different procedures. Thus the procedure is easy to reproduce and a single researcher can prepare multiple different batches of isolated nuclei in a single day, depending on the scale. As a consequence, in one experiment nuclei across multiple experimental conditions (i.e. different genetic backgrounds, different treatment types, and synchronization in different phases of the cell cycle) can be isolated in parallel, which can be further multiplexed due to the ease of cryopreservation of the resulting nuclei. This opens up the study of nuclear structure and function at a throughput that was unthinkable before.

The high speed and ease of preparation not only increases the potential throughput of the method, it also minimizes the handling time, which is key to preserve the structural integrity of the nuclei and to make them usable for functional ex vivo studies. The isolated nuclei are morphologically very well preserved and have largely intact nuclear envelopes that preserve their physiological capacity of excluding fluorescent cargo of a range of sizes. Also nuclear components like the NPC remain structurally intact and retain their ability to bind transport cargo. As case in point, this method has already been used to visualize the nuclear basket of the NPC(Niepel *et al.*), a structure that has been rarely visualized in yeast (Rout and Kilmartin, 1990; Fahrenkrog *et al.*, 1998; Kiseleva *et al.*, 2004), and to show that Mlp1p and Mlp2p are necessary for basket formation and for NE stability(Niepel *et al.*). We expect that the use of this procedure will ease the study of nuclear cytoplasmic transport mechanisms by facilitating the development of cell-free import and export assays considerably facilitating the possibility for multiplexing experimental conditions under study.

## Materials and Methods

### Strains, culture conditions, and spheroplasting

The Nup49p-GFP strain in the BY4741 background was purchased from the Yeast GFP Localization Database (Ghaemmaghami *et al.*; Howson *et al.*). Diploid heterozygous diploid Nup49p-GFP cells were obtained by mating Nup49p-GFP (mat a) with BY4742 (ATCC, accession number 201388).

### Preparation of enriched nuclear fraction

The following volumes are optimized for working with spheroplasts obtained from 1 l yeast cultures. *S. cerevisiae* yeast cells were cultured and spheroplasts were prepared as described (Kipper *et al.*, 2002). All subsequent steps were performed in the cold room at 4°C. Each spheroplast pellet was resuspended in 5 ml of lysis solution [12.5 μl 10% (v/v) Triton X-100 (Surfact-Amps X-100, Pierce, Rockford, IL, USA), 25 μl 1 M DTT, 5 ml polyvinylpyrrolidone (PVP-40; Sigma-Aldrich, St. Louis, MO, USA) solution (8% PVP-40, 20 mM K-phosphate, pH 6.5, 7.5 µM MgCl2].. Immediately before cell lysis 50 μl solution P (20 mg/ml phenylmethanesulfonyl fluoride, 0.4 mg/ml Pepstatin A in ethanol) and 50 μl protease inhibitor cocktail (PIC, P-8340; Sigma-Aldrich, St. Louis, MO, USA) were added. Spheroplasts were lysed mechanically using a Polytron homogenizer (PCU 11 with PTA 10TS Probe; Kinematica, Cincinnati, OH, USA) in intervals of one minute with cooling on ice in between polytroning steps. The polytron was set to 4.5-5 for diploid and 5.5-6 for haploid cells and the probe was completely plunged into the solution. Progress of lysis was checked in between steps by phase contrast microscopy. Approximately 12 ml of 8% PVP-40 solution containing 1:1000 PIC and solution P was added, bringing the volume up to 20 ml and homogenized by a Polytron (Kinematica AG, Luzern, CH) for 30 sec at setting 5.5, fractions pooled and homogenized for another 15 sec. Homogenized cell lysate was loaded on top of a sucrose gradient (8 ml 2.50 M sucrose / BT Mg, 80 μl solution P, 80 μl PIC; 8 ml 1.875 M sucrose / BT Mg, 80 μl solution P, 80 μl PIC) which had been previously prepared in an SW-28 tube (Beckman Coulter, Fullerton, CA, USA) with the boundary between the sucrose layers softened by gentle stirring with a glass rod. Gradients were centrifuged at 28,000 rpm (103,745 g) for 25 min at 4°C in a SW-28 rotor using a Beckman L7-65 ultracentrifuge (Beckman Coulter, Fullerton, CA, USA). After centrifugation nuclei were located at the soft boundary between 1.875 M sucrose and 2.50 M sucrose (see also Fig. 1). The cytosolic fraction in 8% PVP solution, including soluble proteins - and the supernatant fraction, including membranes, mitochondria, vesicles, and microsomes - were carefully unloaded to avoid mixing with nuclei fraction. Optionally, after unloading from the gradient, nuclei were diluted 1:1 in 0.6 M sucrose / BT Mg previously warmed to room temperature. 1:1000 2% DNase I, 1:100 PIC and 1:100 Solution P were added and incubated at room temperature for 20 min with gently inversion every 5 min to create DNase I treated nuclei. 4.0 ml of the resulting nuclear shell suspension were loaded onto a sucrose cushion of 0.4 ml 2.5 M Sucrose / BT Mg and overlaid with 1.0 ml 1.5 M Sucrose/ BT Mg in SW55 tubes and centrifuged using a 55Ti rotor at 45,000 rpm (245,858 x g) for 25 min at 4°C in a L7-65 ultracentrifuge (Beckman Coulter, Fullerton, CA, USA). The nuclei, in approximately 2 M Sucrose / BT Mg, were then dripped into a 50 ml falcon tube filled with liquid nitrogen and the resulting frozen droplets stored at -80μ°C.

### Dyes and staining

The lipid dye DiIC12(3) (1,1'-didodecyl-3,3,3',3'-tetramethylindocarbocyanine perchlorate) was purchased from Molecular Probes (Invitrogen, Paisley, UK), the DNA dye DRAQ5 was purchased from Biostatus (Leicestershire, UK) and R-Phycoerythrin (R-PE) was purchased from Cyanotech Corporation (Kailua-Kona, HI, USA). Bovine serum albumin and ovalbumin were purchased from Sigma-Aldrich (St. Louis, MO, USA). To stain nuclei, the following concentrations and incubation times were used. To stain DNA, DRAQ5 was added to a final concentration of 5 μM. Lipid dye DiIC12(3) was added as a DMSO stock solution to a final concentration of 5 μg/ml without further incubation. Kap95p-Cy5 was incubated with nuclei on ice for 15 min. R-PE, ovalbumin, and bovine serum albumin were added immediately before microscopy. Care was taken for the imaging solutions to maintain at least 1.0 M sucrose so as to not osmotically stress the isolated nuclei. 3.5 μl of the respective nuclei suspension was applied to a glass slide and imaged.

### Fluorescence microscopy

Confocal laser scanning fluorescence microscopy was performed with a Leica TCS SPE system (Leica Microsystems, Wetzlar, Germany) equipped with 488 nm (10 mW output), 532 nm (10 mW output) and 633 nm (18 mW output) and a 63x objective. Epifluorescence illumination combined with differential interference contrast (DIC) microscopy was carried out using a Zeiss Axio Imager Z1 (Jena, Germany) using a 100x objective. Brightness and contrast in images was enhanced using Corel Photo-Paint 12 (Corel Corporation, Ottawa, Kanada) or ImageJ 1.42a (Wayne Rasband, NIH, Bethesda, USA, http://rsb.info.nih.gov/ij/).

### Electron microscopy

Transmission EM was conducted as described (Niepel *et al.*). Fixation steps for scanning EM were performed as described (Kiseleva *et al.*) with minor modifications. Nuclei were immobilized on silicon wafers evaporated with 3 nm chrome and 10 nm gold and subsequently then dried via critical point freezing. After fixation with glutaraldehyde, paraformaldehyde, uranyl acetate, and OsO_4_, the nuclei were evaporated with 2.5 nm PtC. For image acquisition a S-5000 FESEM (Field emission scanning electron microscope) with 30 kV by Hitachi (Hitachi High-Technologies Europe, Krefeld, Germany) was used.

## Author Contributions

All authors collaborated to develop the method and write the manuscript.

## Acknowledgements

We thank Reiner Peters for the supervision and support of JCF during her Ph.D. work, and Brian Chait for laboratory support. Thanks also to Tijana Jovanovic-Talisman for Kap95p-Cy5, and to Patrick Nahirney and Eleana Sphicas of the Bio-Imaging Resource Center at Rockefeller University for help with transmission EM and to Prof. Rudolf Reichelt, Christiane Rasch, and Ulrike Keller at the Institute for Medical Physics and Biophysics, University of Münster for help with SEM. This work was support by NIH grants R01 GM071329, U01 GM098256, and U54 GM103511.

## Additional Files

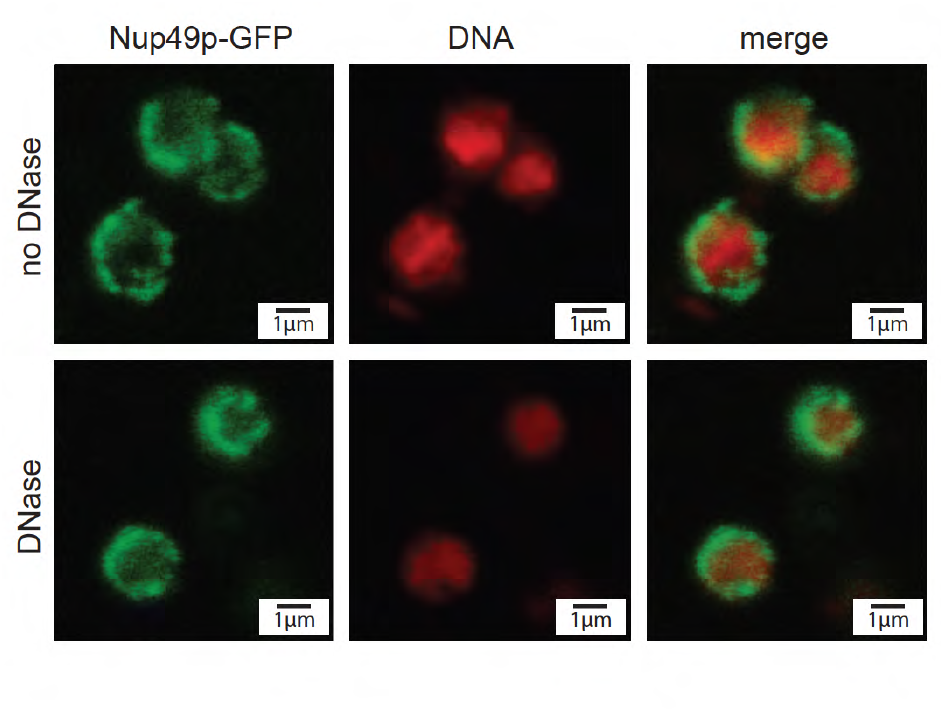

Niepel et al_Figure S1— Supplemental Figure S1— Yeast nuclei and nuclei shells with Nup49-GFP tag, DNA stained with DRAQ5. The panels show fluorescence from Nup49p-GFP (left), DRAQ5-stained DNA (middle), and a merged image (right). Untreated nuclei are shown in the top row and DNase treated nuclei are shown in the bottom. All images have been smoothed using Gaussian blur (radius of 1 pixel).

